# High Throughput Screening with a Primary Human Mucociliary Airway Model Identifies a Small Molecule with Anti-SARS-CoV-2 Activity

**DOI:** 10.1101/2024.05.09.593388

**Authors:** Chandani Sen, Tammy M. Rickabaugh, Arjit Vijey Jeyachandran, Constance Yuen, Maisam Ghannam, Abdo Durra, Adam Aziz, Kristen Castillo, Gustavo Garcia, Arunima Purkayastha, Brandon Han, Felix W. Boulton, Eugene Chekler, Robert Garces, Karen C. Wolff, Laura Riva, Melanie G. Kirkpatrick, Amal Gebara-Lamb, Case W. McNamara, Ulrich A.K. Betz, Vaithilingaraja Arumugaswami, Robert Damoiseaux, Brigitte N. Gomperts

**Affiliations:** UCLA Children’s Discovery and Innovation Institute, Mattel Children’s Hospital UCLA, Department of Pediatrics, David Geffen School of Medicine, UCLA, Los Angeles, CA, 90095, USA; Department of Molecular and Medical Pharmacology, University of California, Los Angeles, CA 90095, USA; California Nanosystems Institute, UCLA, Los Angeles, CA, 90095, USA; Merck KGaA, Darmstadt, Germany; Jonsson Comprehensive Cancer Center, UCLA, Los Angeles, CA, 90095, USA; Eli and Edythe Broad Stem Cell Research Center, UCLA, Los Angeles, CA, 90095, USA; Division of Pulmonary and Critical Care Medicine, Department of Medicine, David Geffen School of Medicine, UCLA, Los Angeles, CA, 90095, USA; EMD Serono Research & Development Institute, Inc., Billerica, MA, 01821, USA; Calibr-Skaggs Institute for Innovative Medicines., 11119 North Torrey Pines Road, La Jolla, CA 92037, USA

**Keywords:** Human mucociliary epithelium, SARS-CoV-2, Respiratory Viral Infections, High throughput drug screening, anti-viral screening, small-molecules, air-liquid-interface, heterogeneity, image quantification, RNA sequencing

## Abstract

Respiratory viruses (e.g. influenza, RSV, SARS etc.) attack the proximal airway and cause a wide spectrum of diseases for which we have limited therapies. To date, a few primary human stem cell-based models of the proximal airway have been reported for drug discovery but scaling them up to a higher throughput platform remains a significant challenge. Here we present a microscale, primary human stem cell-based proximal airway model of SARS-CoV-2 infection, which is amenable to moderate-to-high throughput drug screening. The model recapitulates the heterogeneity of infection seen among different patients and with different SARS-CoV-2 variants. We applied this model to screen 2100 compounds from targeted drug libraries using an image-based quantification method. While there were heterogeneous responses across variants for host factor targeting compounds, the direct acting antivirals showed a consistent response and we characterized a new antiviral drug that is effective against both the parental strain and the Omicron variant.

## INTRODUCTION

The Severe Acute Respiratory Syndrome Corona Virus 2 (SARS-CoV-2) pandemic has resulted in more than 7 million deaths worldwide as of April, 2024[1]. It spurred a global effort to develop new drugs and therapies to combat the virus. Monoclonal antibodies and vaccines were rapidly developed within a year of the pandemic’s start. However, there is still an ongoing search for better small molecules for current and evolving SARS-CoV-2 strains with fewer side effects and drug interactions, especially for immunocompromised and unvaccinated patients. For example, Remdesivir (Veklury) was repurposed for its antiviral effects but requires systemic delivery and the combination drug of the protease inhibitor, nirmatrelvir, together with ritonavir, to reduce its metabolic clearance, is approved under the trade name Paxlovid, and exhibits numerous drug interactions [2]. Molnupiravir (Lageviro) has been shown to drive mutational signatures in global SARS-CoV2 genomes [3]. Efforts to combat the virus have focused on understanding both the host factors [4–7] and viral factors [8–10] influencing the severity of COVID-19.

High throughput drug screening (HTS) of FDA-approved repurposed drug libraries is a preferred strategy for drug discovery because of the compounds’ known safety profile, cost-effectiveness, and time efficiency [11, 12]. There were a lot of drug screening efforts from the onset of the COVID-19 pandemic, using both computational and cell-based *in vitro* methods. Computational methods, powered by artificial intelligence and molecular modeling, have emerged as a valuable ally for COVID-19 drug discovery where deep-learning-based information on drug-target interactions, virtual screening, machine learning, and structure-based drug design have significantly accelerated the identification and optimization of potential therapeutics [13–16]. Although the computational methods accelerate the identification of possible hit compounds, they require further validation by cell-based *in vitro* screening. The traditional *in vitro* high throughput screens (HTS) for respiratory viral infections are often performed on 2D cultures of commercial cell lines like Vero E6, Huh7, Huh7.5, Caco2, Calu-3, A549, and several other non-human cell lines. These cell line-based drug screenings represent an over-simplified, sub-optimal model that has very little or no relevance to the highly specialized human mucociliary airway and its innate immune response.

Air-liquid-interface (ALI) cultures are a well-established model that mimics the micro-physiological conditions of the functional mucociliary epithelium and can also be used to study disease pathogenesis [17–19]. ALI cultures can overcome the limitations of traditional 2D cell-line-based cultures as they demonstrate differentiation to the relevant cell types, epithelial barrier function and an innate immune response. The ALI culture model has been instrumental in demonstrating the mechanisms of SARS-CoV-2 infection of airway cells, the role of exposures (e.g. smoking) on infection severity, and in low throughput drug testing [20–25]. Even though ALI cultures have great potential to study drug efficacy for respiratory viral infections, scaling up the ALI model to a high throughput robust drug screening platform remains a challenge due to the complexity of this culture model, and so, all the prior SARS-CoV-2 studies were done with low throughput. Recently, there has been a growing interest in the development of higher throughput ALI cultures [26, 27] [28], but there are no reports on the optimization of human primary cell-based high throughput ALI systems as a drug screening platform for SARS-CoV-2 infection. Also, there have been no studies to examine the heterogeneity between human airway stem cell donors in their response to SARS-CoV-2 infection in ALI cultures, and there is no information about modeling infection severity across SARS-CoV-2 variants.

Therefore, we developed a 96-transwell primary human mucociliary epithelial ALI culture platform with primary SARS-CoV-2 infection and developed a novel image-based quantification method to quantify this infection. We used primary airway basal stem cells from eight patients with no previous history of lung disease, the parental SARS-CoV-2 strain and three SARS-CoV-2 variants (Beta, Delta, and Omicron), and three drug libraries to screen 2100 repurposed small molecules. We noticed significant heterogeneity of infection based on donors and SARS-CoV-2 variants. We also found a lack of consistency of efficacy of drugs targeting host factors. However, when screening with direct-acting antivirals, we found consistent results across all donors and SARS-CoV-2 variants. We used our 96-transwell ALI platform to characterize a new small molecule with antiviral properties, which showed efficacy against the parental strain and the Omicron variant. This 96-transwell primary human mucociliary epithelial ALI culture platform with SARS-CoV-2 infection shows potential for drug screening for respiratory viral infections.

## RESULTS

### 96-transwell air-liquid-interface (96-TW ALI) model of human mucociliary airway for drug screening demonstrates heterogeneity of infection among different airway donors and SARS-CoV-2 variants

We previously showed that live SARS-CoV-2 virus can infect the 24-transwell ALI (24-TW ALI) model of human mucociliary epithelium [21]. Since the 24-TW ALI model is sub-optimal for high throughput screening, we optimized a 96-TW ALI for screening purposes. The detailed protocol of 96-TW ALI generation is described in the STAR methods section and the general timeline is shown in Figure 1A. Briefly, we isolated airway basal stem cells (ABSC) from the trachea of donors with no history of prior lung disease, and the cells were seeded on pre-collagen-coated transwells. Once the cells formed a confluent monolayer (∼5 days), media was removed from the top chamber and the surface of the cells was exposed to air, and the ALI condition was initiated. After 21 days of ALI culture, when they closely resemble the airway epithelial barrier of the cartilaginous airways and are well differentiated with ciliated, mucus, and basal cells and an intact innate immune response, we infected them with SARS-CoV-2 virus.

**Figure 1:**
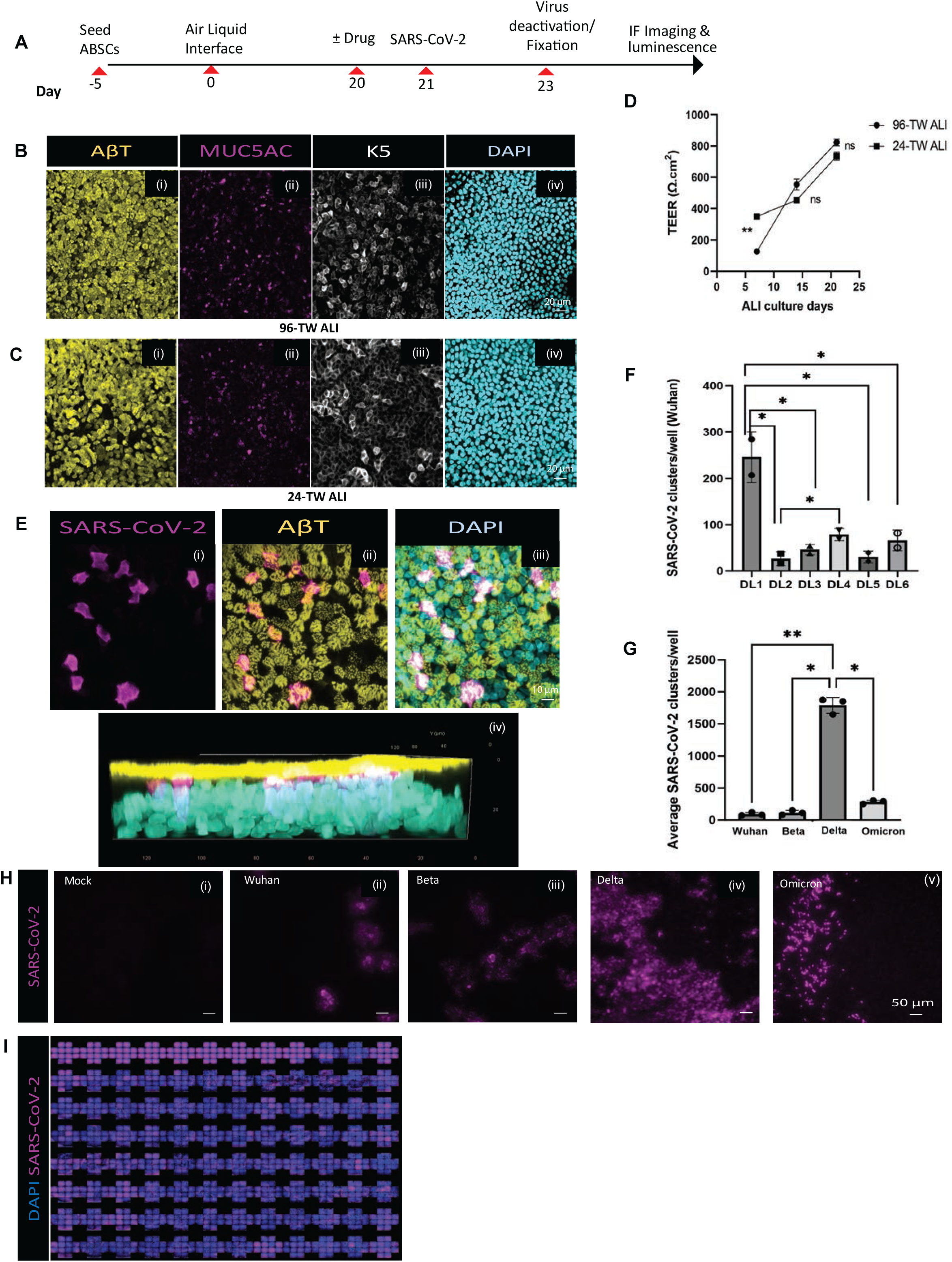
Development of a human primary mucociliary airway stem cell-based SARS-CoV-2 infection model. **A.** Schematic of the experimental timeline. **B.** 96-TW ALI and **C.** 24-TW ALI with immunofluorescent (IF) staining showing **(i)** Nuclei (DAPI), **(ii)** Ciliated cells (AβT), **(iii)** Mucus cells (MUC5AC), **(iv)** Basal stem cells (K5). Scale bar = 50uM. **D.** TEER values of 24-and 96-TW ALI. P values are calculated from technical replicates using Student’s t test. **: P<0.01 **E.** Confocal imaging (magnification: 63X) of IF staining showing a typical SARS-CoV-2 (Omicron) cluster of infection in the 96-TW ALI model. **(i)** SARS-CoV-2 infection (Spike)**. (ii)** Ciliated cells (AβT) with SARS-CoV-2. **(iii)** Ciliated cells (AβT) with SARS-CoV-2 and DAPI. **(iv)** Orthogonal projection of IF staining of the infected 96-TW ALI culture. **F.** Quantification of degree of infection among different human donors. DL=Donor Lung samples. P values are calculated from technical replicates using one way ANOVA test. *: P<0.05. **G.** Quantification of the average degree of infection among different SARS-CoV-2 Variants. P values are calculated from technical replicates using one way ANOVA test. **: P<0.01, *: P<0.05. **H.** Variability of degree of infection by IF staining for SARS-CoV-2 in ii) Wuhan (Spike antibody), iii) Beta (Spike antibody), iv) Delta (Nucleocapsid antibody), v) Omicron (Nucleocapsid antibody). Scale bar = 50uM. **I.** Representative HTS images of SARS-CoV-2 infection (Nucleocapsid antibody) with 96-TW ALI model.

We compared the 96-TW ALI (Figure 1B) with the current gold standard 24-TW ALI (Figure 1C) and found 96-TW ALI cultures closely resembled the airway epithelial barrier of the mucociliary airways and are well differentiated with (i) ciliated cells, (ii) mucus cells (iii) basal cells and (iv) uniform nuclei distribution, and have no difference in distribution of these cells when compared with the 24-TW ALI (Figure 1B,C, i-iv). We also measured trans-epithelial electrical resistance (TEER) as a measure of tight junction formation of the ALI epithelial barrier, and the week-wise values for both 24-TW- and 96-TW-ALI were within the acceptable range and not significantly different at days 15 and 25 of culture (Figure 1D). All these data suggest that the 96-TW human ALI model is a relevant model of the mucociliary epithelium and can replace the 24-TW human ALI model to achieve higher throughput.

We identified some key factors for generating a successful image-grade 96-TW ALI with uniform infection quality. Firstly, the transwell membrane material is important for imaging. Although human ABSCs grow well on both polycarbonate (PC) and polyester (PET) membranes, only PET membranes are transparent, which allows much higher quality image acquisition during high throughput confocal microscopy. Figure 1E (i-iv) demonstrates a typical cluster of SARS-CoV-2 infection in ciliated cells in our PET based 96-TW ALI that proves its suitability for high-content imaging. Second, we found that the number of cells seeded is important. We observed that the optimal cell density to plate cells is a function of active ABSCs in the isolated population. So, expanding the isolated ABSCs up to passage 1 (P1) results in activation and enrichment of the ABSC population and requires less initial cell numbers to obtain a confluent monolayer (10,000 cells/well for unexpanded cells, 8,000 cells/well for expanded), and less time to reach confluency of ABSCs (7 days for unexpanded, 5 days for expanded). Apart from cell density and submerged culture duration, other quality parameters were unaltered between P0 and P1, whereas expanding the cell beyond P2 resulted in poor culture quality with uneven cell type distribution and absence of tight junction formation due to reduction in ABSC numbers. Thirdly, we identified donor heterogeneity and the need to pool donor cells as being key for reproducible drug screening. For this study we used eight airway donor lung (DL) samples, and we noticed a large degree of variation in the percentage of infected cells from donor to donor (Figure 1F). The variation seen in SARS-CoV-2 infection of ALI cultures from different donors is likely due to differences in the host innate immune response. Even though recapitulation of this heterogeneity of infection in our model reveals its suitability for personalized medicine, for the purposes of finding a hit compound that has efficacy for a large patient population, we need to have a reliable, reproducible response. Therefore, we pooled ABSCs from three different patients to generate ALI cultures. We followed hepatocyte studies where pooling of cells from different patients is standard practice in drug screening [29]. The demographics of all the ABSC donors used for this study are elucidated in Supplementary Table 1.

Another crucial parameter for high throughput drug screening is a fast readout with quantification. For our high-content screening, we scanned across 14 sites in each well of the 96-TW ALI and built an adaptive thresholding-based algorithm to quantify infection based on cluster properties like number and intensity across all sites. For this work, we selected infection cluster count/well as our preferred read-out as it was more consistent and convenient to validate with ImageJ-based counting. The detailed methods for the quantification are provided in the methods section.

Once the screen was developed and validated for SARS-CoV-2 infection, we initially infected the ALI cultures with the Wuhan parental SARS-CoV-2 strain (Isolate USA-WA1/2020, BEI Resources NR-52281) at a low MOI (0.1), followed by the Beta (Isolate hCoV-19/USA/MD-HP01542/2021, BEI Resources NR-55282), Delta (Isolate hCoV-19/USA/MD-HP05647/2021 BEI Resources NR-55672), and Omicron (Isolate hCoV-19/USA/MD-HP20874/2021, BEI Resources NR-56461) variants as they arose during the pandemic. Figure 1G and H (i-v) demonstrate the variability of infection among these strains in the 96-TW ALI and the corresponding quantification. The mock (no infection) well had no interfering signal or noise (Figure 1G,Hi). The parental strain had the lowest level of infected clusters per well of 96-TW ALI (99 +/- 15) (Figure 1G,Hii), while the Beta variant had an average of 120+/-21 (Figure 1G,Hiii). Delta showed the highest level of infectivity with an average number of 1876+/-102 infection clusters (Figure 1G,Hiv), while the Omicron variant showed 280+/-23 infected clusters per well (Figure 1G,Hv). The different degree of infectivity across the SARS-CoV-2 variants appeared to be associated with the known degree of disease severity of the variants. We noted differences across viral variants with some having many small sized infection clusters suggesting that for that variant, entry is easier than replication. And for other variants, there were large clusters of infection that were few in numbers suggesting entry was difficult, but replication was easier for that variant. The SARS-CoV-2 infected ALI cultures are therefore a potential model to predict response to therapies that are currently in the clinic and to screen for novel compounds. Therefore, we next tested our model for drug screening in HTS format (Figure 1I).

### Drug discovery using the 96-TW ALI model of SARS-CoV-2 infection

To validate our 96-TW ALI drug screening platform, we calculated the Z-factor (Z’) [30] of the 96-TW ALI model to test its efficacy for high throughput drug screening (HTS) for SARS-CoV-2 from mock (no virus) and infection (virus only, no drug) based on cluster count. The calculated average Z’ for each of the infections (Wuhan 0.967, Delta 0.995, Omicron 0.614) proved its suitability to be used for screening for novel drugs for SARS-CoV-2 (Figure 2A). Beta, however, did not show any significant Z’ (data not shown). For rapid drug screening and quantification purposes, in parallel to the image-based quantification, we also tested and validated the SARS-CoV-2 Wuhan virus reporter NanoLuc Luciferase (Nano-Glo Luciferase assay, Promega) in our system (average Z’ Nanoluc 0.588) (Figure 2A).

**Figure 2:**
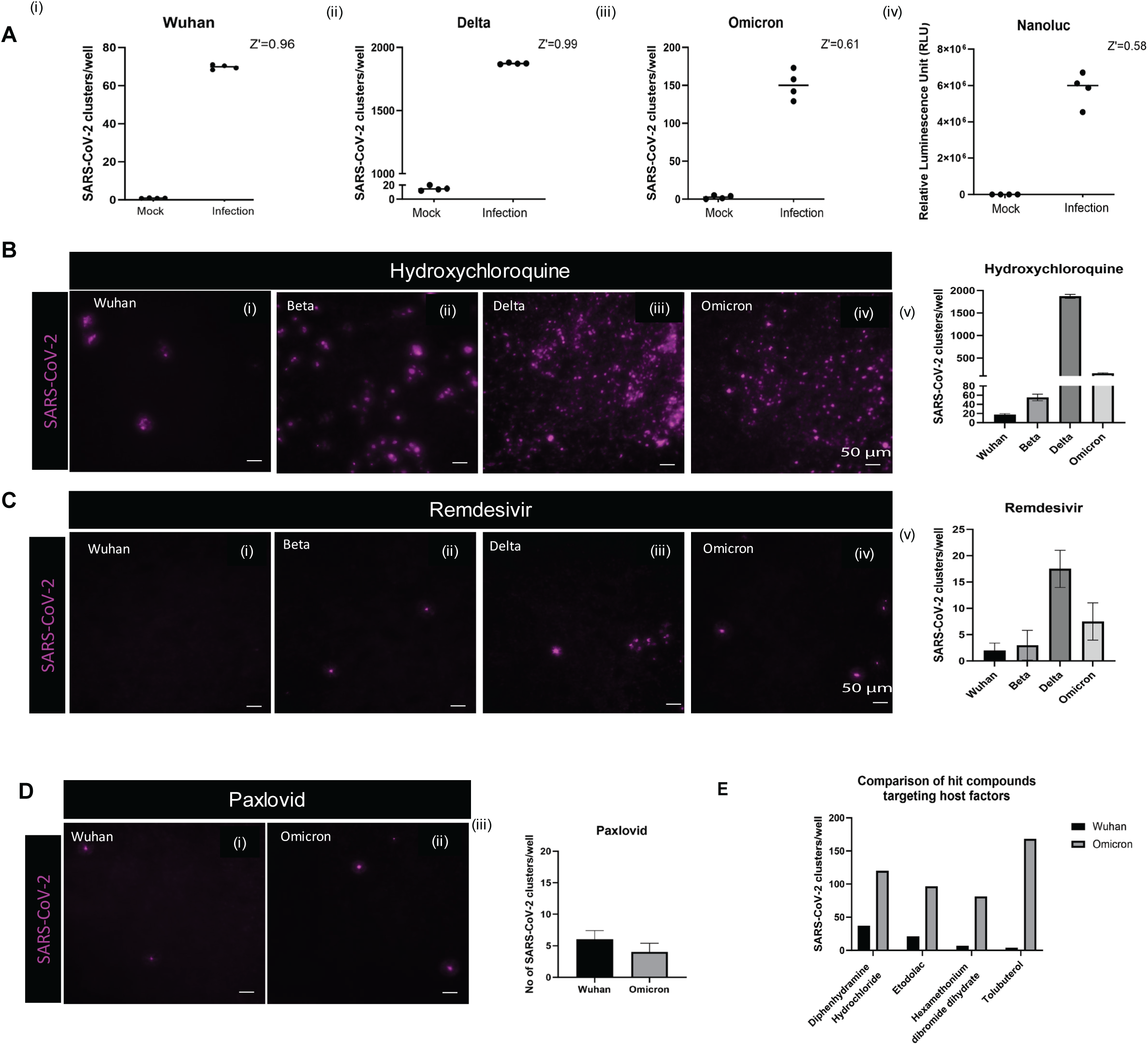
Validation and testing of the 96-TW as drug screening platform. **A.** Z’ values across successive SARS-CoV-2 Strain infections: (i) Wuhan, (ii) Delta, (iii) Omicron, and in the (iv) Wuhan-Nanoluciferase (Nanoluc) reporter assay. **B.** IF staining showing efficacy of Hydroxychloroquine across different strains: (i) Wuhan (Spike antibody), ii) Beta (Spike antibody), iii) Delta (Nucleocapsid antibody), iv) Omicron (Nucleocapsid antibody). (v) shows quantification of Hydroxychloroquine efficacy. Scale bar = 50uM. **C.** IF staining showing efficacy of Remdesivir across different strains: (i) Wuhan (Spike antibody), ii) Beta (Spike antibody), iii) Delta (Nucleocapsid antibody), iv) Omicron (Nucleocapsid antibody). (v) shows quantification of Remdesivir efficacy. Scale bar = 50uM. **D.** Efficacy of Paxlovid against (i) Wuhan (Spike antibody), ii) Omicron (Nucleocapsid antibody). (iii) Quantification of Paxlovid efficacy. Scale bar = 50uM. **E.** Top hits from the drug screen libraries detailed in Supplementary Table 1, showing differences in efficacy in 96-TW ALI cultures infected with Wuhan versus Omicron SARS-CoV-2 variants.

We examined the response of the SARS-CoV-2 parental strain and variants in 96-TW ALI to hydroxychloroquine (10µM) and found that this efficacy was reduced across each successive variant (Figure 2Bi-iv) which we quantified in Figure 2Bv (Wuhan:17.5±1.5,Beta: 55±5, Delta:1870.5±31.5, Omicron:160±10). Unlike hydroxychloroquine, we found a consistent response of our 96-TW ALI cultures infected with all SARS-CoV-2 variants to Remdesivir (10µM) (Figure 2Ci-iv) with quantification in Figure 2Cv (Wuhan:2±.5, Beta:3±1, Delta:17.5±1.25, Omicron:7.5±1.25). We also found that Paxlovid (Nirmatrelvir-Ritonavir) was effective against the parental strain and the Omicron variant in our 96-TW ALI model (Wuhan:6±.5, Omicron: 4±.5)) (Figure 2Di-iii).

For drug screening in 96-TW ALI, we tested 2100 compounds across 3 libraries [440 from LOPAC (Sigma Aldrich), 660 from New Prestwick (Greenpharma), 1000 from NKIL (Selleck Chemicals)] in 8 airway donor lung (DL) samples, against Wuhan, Beta and Delta and found a total of 72 compounds that reduced infection, elucidating a 2.8% hit rate. Notably, we included 13 NKIL hits that we previously identified when screening with Vero cells [22, 30] and they were not effective in the human 96-TW ALI model. We also found that the hit compounds were different for different variants indicating that the inhibition mechanism for each variant may be different. All 72 hit compounds from our primary screen were re-screened against Wuhan in one 96-TW ALI culture, and only 4 compounds – Diphenhydramine, Etodolac, Hexamethonium dibromide dihydrate, and Tulobutarol were effective (average no of COVID-19 clusters/well: 13.33) as they reduced infection by ∼7 fold compared to no drug control (99 clusters/ well) (Figure 2E). However, when tested against Omicron they were not nearly as effective (average no of COVID-19 clusters/well: 116.66) (Figure 2E). Our 96-TW ALI drug screen did not identify any compounds that were effective against the Wuhan parental strain and the Omicron variant. But the most effective drugs in our assay were Paxlovid and Remdesivir, which are both direct acting antivirals. We therefore collaborated with Merck KgaA, Darmstadt, Germany to test some of their direct-acting antivirals in our 96-TW ALI screening platform.

### Screening with 96-TW ALI cultures exhibits a new compound with antiviral defense properties

In collaboration with Merck KgaA, Darmstadt, Germany, we tested a custom library of small molecules identified from antiviral chemical space in our 96-TW ALI cultures and singled out one small molecule (Merck KgaA, Darmstadt, Germany Compound#49, abbreviated as MKGaA#49 henceforth) that showed a significant reduction in infection against the parental strain (Figure 3Ai) (average no of COVID-19 clusters/well: 3) and the Omicron variant (Figure 3Aii) (average no of COVID-19 clusters/well: 5)(Fig 3Aiii). When compared with Remdesivir, MKGaA#49 showed comparable efficacy against Omicron (average no of COVID-19 clusters/well for Remdesivir: 2.4) (Figure 3Bi) and in Wuhan Nanoluc reporter assay (Figure 3Bii).

**Figure 3:**
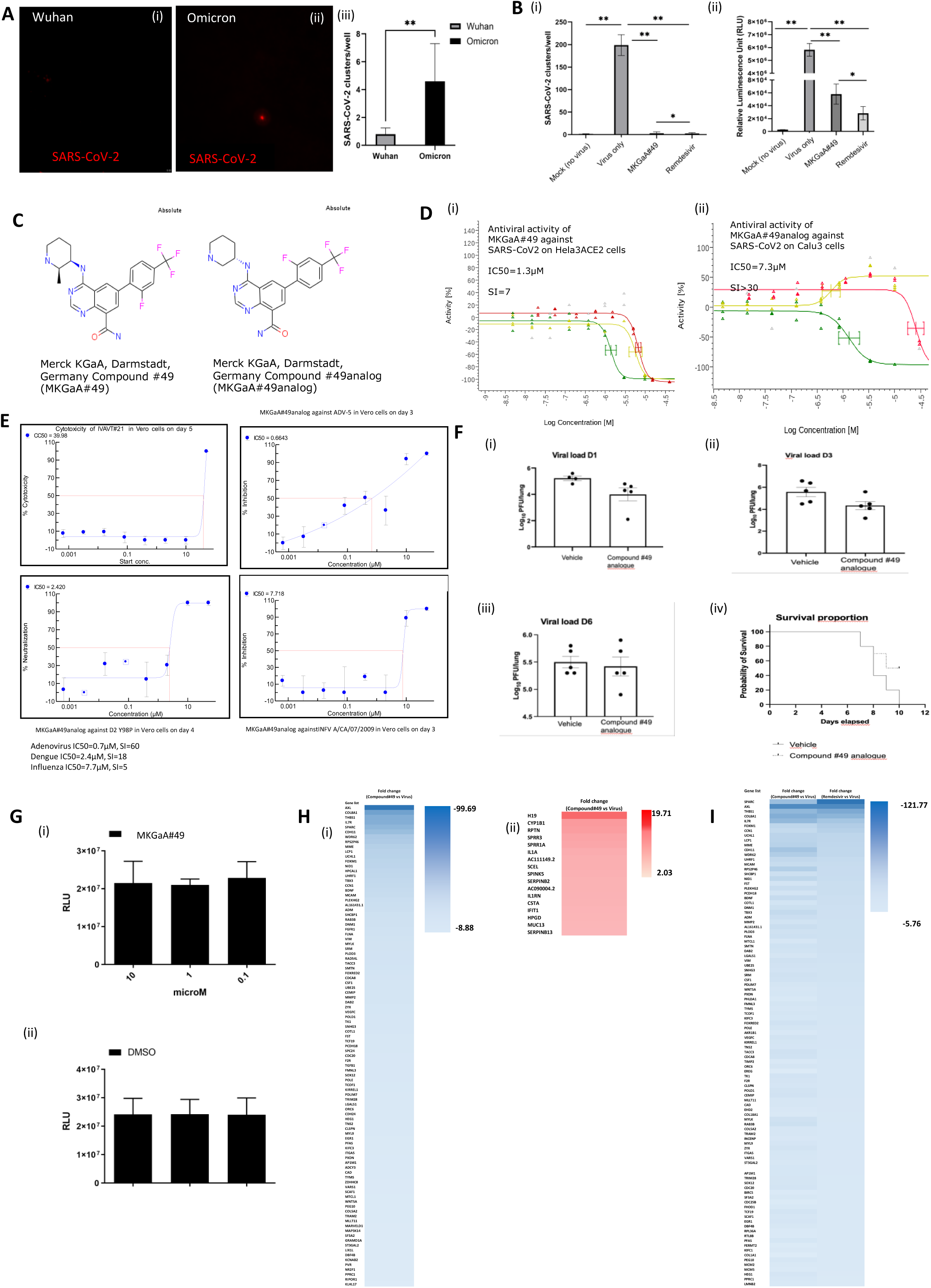
The screen identifies a compound with antiviral efficacy against SARS-CoV-2. **A.** IF staining showing the efficacy of MKGaA#49 (Merck KGaA, Darmstadt, Germany Compound#49)against (i) Wuhan and (ii) Omicron infection in the 96-TW ALI cultures. Scale bar = 50uM. (iii) Quantification of MKGaA#49 efficacy. **B.** Comparative efficacy of MKGaA#49 and Remdesivir in reducing Omicron infection by (i) IF image quantification (nucleocapsid antibody), and (ii) Nanoluc assay. **C.** Structure of MKGaA#49 and a MKGaA#49analog. **D.** Antiviral activity in Hela3ACE2 cells (i) and Calu3 cells (ii). Graphs show antiviral activity measured with a SARS-CoV-2 immunofluorescence signal leading to identification of infected cells with 0% activity equals 100% infected cells (green curve), total cells per well in SARS-CoV-2 infected cell test with 0% activity equaling no change vs. control (yellow curve), total cells per well in HeLa-ACE2 uninfected cell control (red curve). **E.** Antiviral activity of MKGaA#49 against Adenovirus, Dengue and Influenza. **F.** *In vivo* antiviral efficacy of MKGaA#49 analog. Mice exhibited reduced viral load at Day 1 post infection (i) and Day 3 post infection (ii) although there was no difference seen in viral load at Day 6 post infection (iii). Mice exhibited increased survival upon drug treatment (iv). **G.** Cell Titre Glo assay to assess for any toxicity of (i) MKGaA#49 across three concentrations as compared to (ii) DMSO alone across the same concentrations. **H.** Global RNA sequencing of SARS-CoV-2 infection compared to SARS-CoV-2 infection with MKGaA#49. Heat map of (i) top 100 downregulated genes and (ii) all 17 upregulated genes. **I.** Global RNA sequencing of SARS-CoV-2 infection compared to SARS-CoV-2 infection with MKGaA#49 or Remdesivir. Heat map of top 100 genes differentially expressed in SARS-CoV-2 infection alone versus SARS-CoV-2 infection treated with MKGaA#49 compared to the top 100 genes differentially expressed after SARS-CoV-2 infection treated with Remdesivir versus SARS-CoV-2 virus alone.

The structure of MKGaA#49 (6-(2-Fluoro-4-trifluoromethyl-phenyl)-4-((2R,3R)-2-methyl-piperidin-3-ylamino)-quinazoline-8-carboxylic acid amide) and its close analog:MKGaA#49analog (6-(2-Fluoro-4-trifluoromethyl-phenyl)-4-((2R,3R)-2-methyl- piperidin-3-ylamino)-quinazoline-8-carboxylic acid amide) are shown in Figure 3C. MKGaA#49 had previously been identified to show antiviral properties against the SARS-CoV-2 parental strain on Hela3ACE2 cells (Figure 3Di) and on Calu3 cells (not shown) with an IC50 of 1.3µM and 7µM respectively. The MKGaA#49analog was also tested for antiviral activity against the SARS-CoV- 2 parental strain in Hela3ACE2 cells (not shown) and in Calu3 cells (Figure 3Dii) and observed strong antiviral activity with an IC50 of 2.4µM and 7.3µM respectively. The MKGaA#49-analog was also tested for activity against a series of viruses and exhibited antiviral activity against Adenovirus (IC50=0.7µM), Dengue (IC50=2.4µM) and Influenza (IC50=7.7µM) as shown in Figure 3E. No activity was observed against HSV-1, HSV-2, Chikungunya and Zika (data not shown).

We also evaluated MKGaA#49-analog for *in vivo* activity in a mouse model. As shown in Figure 3F, oral treatment of mice with the compound resulted in reduced viral titers in the lungs on Day 1 (D1) (Fig 3Fi), D3 (3Fii) and D6 (Fig 3Fiii). Likewise, a prolonged survival could be observed in mice treated with the compound compared to vehicle control (Fig 3Fiv). When tested with the

Cell Titre Glo assay, MKGaA#49 showed very low toxicity, and the results were essentially the same as the solvent DMSO alone (Figure 3Gi,ii).

Given the compound’s efficacy both *in vitro* and *in vivo*, next we sought to understand its mechanism of action and we therefore assessed the effect of MKGaA#49 treatment compared to DMSO treatment in SARS-CoV-2 infected 96-TW ALI cultures by global RNA sequencing. We found many genes that were downregulated by MKGaA#49 in infected ALI cultures and only about 50 genes that were upregulated (Figure 3H). Comparison of MKGaA#49 with Remdesivir showed similar gene expression changes (Figure 3I). Analysis of the GO terms for these differentially expressed genes showed the antiviral response, and innate immune response were induced by MKGaA#49, in addition to negative regulation of viral genome replication (Table 1).

**Table 1.** Functional classification of differentially expressed genes between MKGaA#49 versus DMSO treated ALI cultures after SARS-CoV-2 Virus infection (Wuhan variant)

| **Annotation Cluster 1** | **Enrichment Score:**<br>**9.7** | **Count** | **P_Values** | **Benjamini** |
| --- | --- | --- | --- | --- |
| UP_KW_BIOLOGICAL_PROCESS | Antiviral defense | 13 | 1.10E-15 | 2.70E-14 |
| GOTERM_BP_DIRECT | defense response to virus | 14 | 2.00E-15 | 7.70E-13 |
| GOTERM_BP_DIRECT | response to virus | 11 | 3.80E-14 | 7.50E-12 |
| GOTERM_BP_DIRECT | negative regulation of viral genome replication | 9 | 7.30E-14 | 9.60E-12 |
| UP_KW_BIOLOGICAL_PROCESS | Innate immunity | 13 | 4.50E-10 | 5.60E-09 |
| UP_KW_BIOLOGICAL_PROCESS | Immunity | 14 | 5.30E-07 | 4.40E-06 |
| GOTERM_BP_DIRECT | innate immune response | 10 | 6.90E-06 | 5.40E-04 |
| GOTERM_MF_DIRECT | RNA binding | 6 | 2.80E-01 | 1.00E+00 |
| **Annotation Cluster 2** | **Enrichment Score:**<br>**5.63** | **Count** | **P_Values** | **Benjamini** |
| KEGG_PATHWAY | Influenza A | 8 | 2.00E-07 | 1.60E-05 |
| KEGG_PATHWAY | Hepatitis C | 7 | 2.60E-06 | 1.00E-04 |
| KEGG_PATHWAY | Coronavirus disease - COVID-19 | 7 | 2.40E-05 | 6.40E-04 |
| **Annotation Cluster 3** | **Enrichment Score:**<br>**3.72** | **Count** | **P_Values** | **Benjamini** |
| GOTERM_CC_DIRECT | cornified envelope | 7 | 4.10E-09 | 2.70E-07 |
| GOTERM_BP_DIRECT | keratinocyte differentiation | 6 | 3.20E-07 | 3.20E-05 |
| INTERPRO | Cornifin (SPRR1) | 3 | 1.50E-05 | 1.70E-03 |
| GOTERM_BP_DIRECT | epidermis development | 5 | 3.70E-05 | 2.20E-03 |
| GOTERM_BP_DIRECT | peptide cross-linking | 4 | 4.00E-05 | 2.20E-03 |
| GOTERM_BP_DIRECT | keratinization | 5 | 4.70E-05 | 2.30E-03 |

## DISCUSSION

Here we have developed a primary human airway cell based 96-TW ALI model as a drug screening platform for SARS-CoV-2 variants. The 96-TW ALI recapitulated the heterogeneity of infection that is seen clinically among people infected with SARS-CoV-2 and also across infections with different SARS-CoV-2 variants. 96-TW ALI drug screening of FDA approved and re-purposed drug libraries targeting host factors across variants did not reveal any consistent hits. However, direct-acting antivirals like Remdesivir and Paxlovid were consistently effective in our model against multiple SARS-CoV-2 variants. When screened against a panel of antivirals from Merck KGaA, Darmstadt, Germany, our model identified a novel compound (MKGaA#49) that showed similar efficacy to Remdesivir against the SARS-CoV-2 parental strain (Wuhan) and the Omicron variant. It also showed gene expression differences suggesting antiviral defense, induction of the innate immune response and inhibition of viral genome replication as its mechanism of action.

Overall, this study highlights a relevant, novel, human drug screening platform for SARS-CoV-2 drug discovery. The study also demonstrates how donor diversity and SARS-CoV-2 variants play a key role in therapeutic discovery and how pooling donors can improve the variability of the assay. Our 96-TW ALI model also has potential to be used for drug discovery for other respiratory viral infections. An important aspect of our model is the fact that it has innate immunity and therefore more closely recapitulates respiratory viral infections of the human airway. While ALI cultures have been used for decades, they have not been used for moderate to high throughput drug screening for viral infections because throughput has been lacking. This is because high throughput imaging and quantification have been difficult to achieve. We have overcome these barriers and, in the process, identified MKGaA#49, which shows potential for treating multiple variants of COVID-19. We also speculate that it could possibly be used to treat other respiratory viruses. Our data also demonstrates that compounds targeting host factors from FDA-approved and repurposed drug libraries show variability in response across donors while direct acting antivirals were effective against all donors and variants. While pooling donors reduced variability, it also showed a lack of efficacy of the compounds targeting host factors. This suggests that the direct acting antiviral chemical space should be expanded and mined with assays such as ours to identify the next generation of therapies for respiratory viral infections and possible future pandemics.

### Limitations of the study

While we speculate that the throughput could be advanced even further on our 96-TW ALI culture model, we were only able to achieve moderate throughput with our available technology. We speculate that as high throughput confocal imaging technologies develop, our 96-TW ALI culture model will become higher throughput and the quantification algorithms will become more sophisticated. We were also limited in the age range of airway donor tissue we received, so we did not study variability in infection in young children as compared to older adults.

## ACKNOWLEDGEMENTS

This work was supported by Merck KGaA, Darmstadt, Germany, Bill & Melinda Gates Foundation, CIRM DISC2 19-11764 award (BG), the NIH/NCI Grant R01CA208303 (BG), the Department of Defense PR202868 award (BG), Mercatus Center Fast Grants for COVID-19 research (BG), the UCOP Emergency Funding COVID19 TRDRP Seed grant Award R00RG2383 (BG), the UCLA Oversight COVID-19 Research Committee (OCRC)(BG and VA), the UCLA Eli & Edythe Broad Center of Stem Cell Research (BSCRC) (BG).

We greatly appreciate the UCLA BSCRC Microscopy Core, and the UCLA Translational Pathology Core Laboratory. This research was supported by NIH National Center for Advancing Translational Science (NCATS) UCLA CTSI Grant Number UL1TR001881. The following reagents were obtained through BEI Resources, NIAID, and NIH: Polyclonal Anti-SARS Coronavirus (antiserum, Guinea Pig), NR-10361, and Nucleocapsid Ab (GeneTex, GTX135357). We are grateful to Dirk Finsinger, Markus B Klein, Sven Poetzsch, Anna Coenen-Stass, Lukas Friedrich, Jörg Zissel, Jan-Carsten Pieck, Momar Toure, Constantin Neagu, James Cummings, Bin Li and Andrea Unzue-Lopez, Theresa Johnson.

## AUTHOR CONTRIBUTIONS

**C.S.:** conception and design, collection and assembly of data, data analysis and interpretation, manuscript writing,

**G.G. J., T.M.R., A.V.J., A.P., A.D., A.A., C.Y., M.G., K.C., B.H., F.W.B. E.C., R.G., K.C.W., L.R., M.G.K., A.G-L., C.W.M:** data analysis and interpretation and technical support.

**V.A., R.D., U.B.:** conception and design, data analysis and interpretation, manuscript writing, financial support

**B.N.G.:** conception and design, data analysis and interpretation, manuscript writing, final approval of the manuscript, and financial support.

## RESOURCE AVAILABILITY

### Lead contact

Further information and requests for resources and reagents should be directed to and will be fulfilled by Lead Contact, Brigitte N. Gomperts.

### Materials Availability

This study did not generate new unique reagents.

### Data and Code Availability

N/A

## METHODS

### Experimental Models and Subject Details Human Tissue Procurement

Large airways and bronchial tissues were acquired from three different tissue sources: 1. de-identified normal human donors after lung transplantations at the Ronald Reagan UCLA Medical Center. Tissues were procured under an Institutional Review Board-approved protocol at the David Geffen School of Medicine at UCLA (IRB#21-000390-CR-00003). 2. Normal human bronchial epithelial cells (NHBE) from non-smokers were obtained from Lonza and all samples were de-identified. 3. IIAM donor lung and trachea samples were obtained with institutional approval. De-identified patient information for all samples is shown in Supplementary Table 1.

### ABSC Isolation

Human ABSCs were isolated following a previously published method from our laboratory [21, 31–33]. All steps were performed having the trachea in cold PBS with antimicrobials. Briefly, proximal cartilaginous airways were dissected, cleaned, and incubated in 50U/mL dispase for 45 minutes at room temperature. Tissues were then incubated in 0.5mg/mL DNase for another 45 minutes at room temperature. Epithelium was stripped and scraped with a cell-scraper and incubated in 0.25% Trypsin-EDTA for 30 minutes shaking at 37°C to generate a single cell suspension. Isolated cells were passed through a series of strainers, 500 μm, 300 μm, and 100 μm strainer and either plated directly (Passage 0 (P0)) for Air Liquid Interface (ALI) cultures or expanded in collagen IV (0.5mg/mL) coated T25 or T75 up to P1/P2 and then plated for ALI cultures.

### 96-TW ALI Cultures

96-TW with 0.4μm pore polyester membrane inserts (Corning 7369) were coated with 0.5mg/mL collagen type IV and allowed to air dry overnight inside a biosafety cabinet. ABSCs (P0-P2) were seeded at 8000-10,000 cells per well onto collagen-coated transwells in Pneumacult Ex Plus media (STEMCELL Technologies), supplemented with Primocin (Invivogen), 1X Penicillin-Streptomycin-Neomycin (PSN) Antibiotic Mixture (Thermo Fisher Scientific). ABSCs were allowed to grow in the submerged phase with 180μl media in the basal chamber and 75μl media in the apical chamber until they were 80-100% confluent and tight cell junctions were formed (5- 7 days). ALI cultures were then established via airlifting, and cells were cultured with 180μl Pneumacult ALI (STEMCELL technologies) media only in the basal chamber, for 21 days until viral infection. Media was changed every other day and cultures were maintained at 37°C and 5% CO2. In the case of drug testing, drugs were added at their required concentration in the basal chamber media 24 hours before viral infection.

### SARS-CoV-2 infection

SARS-CoV-2 strains including Isolate USA-WA1/2020 (Parental, BEI Resources NR-52281) at a low MOI (0.1), followed by the Isolate hCoV-19/USA/MD-HP01542/2021 (Beta, BEI Resources NR-55282), Isolate hCoV-19/USA/MD-HP05647/2021 (Delta, BEI Resources NR-55672), and Isolate hCoV-19/USA/MD-HP20874/2021 (Omicron, BEI Resources NR-56461) were obtained from Biodefense and Emerging Infectious (BEI) Resources of the National Institute of Allergy and Infectious Diseases (NIAID). All the studies involving live viruses were conducted in the UCLA BSL3 high-containment facility with appropriate institutional biosafety approvals. SARS-CoV-2 was passaged once in Vero-E6 cells and viral stocks were aliquoted and stored at -80°C. Virus titer was measured in Vero-E6 cells by TCID_50_ assay [22, 34].

ALI cultures on the apical chamber of transwell inserts were infected with SARS-CoV-2 viral inoculum (MOI of 0.1; 100 µl/well) prepared in PneumaCult ALI (STEMCELL technologies) media. The basal chamber of the transwell contained 180 µl of ALI media. For mock infection, ALI media (30 µl/well) alone was added. The inoculated plates were incubated for 2 hr at 37 °C with 5% CO2. At the end of incubation, the inoculum was removed from the apical chamber. At selected time points live cell images were obtained by bright field microscopy to detect cytopathic effect (CPE) in SARS-CoV-2 infected cells, indicating viral replication and associated cell injury. At 72 hours post-infection (hpi), ALI cultures were fixed in 4% paraformaldehyde followed by immunofluorescence (IF) analysis.

### NanolucIferase

We performed Nano-Glo Luciferase Assay System as indicated by the manufacturer for assessing the presence of luciferase activity. ALI Cultures were pretreated with drug compounds for 24 hours. Pretreated ALI cultures were then infected with SARS-Related Coronavirus 2, Isolate USA- WA1/2020 (icSARS-CoV-2-nLuc) (BEI Resources NR-54003) After 48 hpi a working reagent luciferin (100 μL) was added to the cells and incubated for at least 3 min at room temperature. Thereafter, the luminescence of each condition was measured in triplicate values and recorded. Percent viability for each compound was calculated based on vehicle (water or DMSO) treated cells.

### Immunofluorescence and Confocal Imaging

Post fixation with 4% PFA, the ALI cultures were washed 3 times with Tris-Buffered Saline and Tween-20 (TBST) followed by permeabilization with 0.5% Triton-X for 10 minutes. ALIs were then blocked using serum-free protein block (Dako X090930) for one hour at room temperature and overnight for primary antibody incubation. Based on preliminary trials, we optimized different SARS-CoV-2 antibodies for different strains of SARS-CoV-2 virus. SARS-CoV [BEI Resources: NR-10361 Polyclonal Anti-SARS Coronavirus (antiserum, Guinea Pig)] for Wuhan and Beta strains; SARS-CoV-2 [(2019-nCoV) Spike S1 Antibody, Rabbit Mab] for Delta strain, and SARS-CoV-2 nucleocapsid antibody (GeneTex, GTX135357) for the Omicron strain. The full list of antibodies used in this study is provided in the Key Resources Table as part of the STAR Methods. After several washes with TBST, secondary antibodies were incubated on samples for 1 hour in darkness, washed, and used for confocal imaging. High-throughput, high-content confocal images were obtained using ImageXpress (Molecular Devices) with a 20X objective and number of infection clusters per well were quantified by developed algorithm. To compare ALI differentiation in 24- and 96-TW (Figure 1), images were captured on Zeiss LSM 880 confocal microscopes.

### Image Quantification

Plates were imaged using a Molecular Devices Confocal Imager at 20x resolution with an extra long working distance (ELWD) objective (n/a =0.45) with at least 14 fields per well on an Image Xpress Confocal (Molecular Devices) while avoiding the corners of the well. Each site in each well was individually focused by images for the DAPI channel and pictures obtained for both the DAPI and TRITC channel. Images were streamed real time into a SQL managed instance of MetaXpress Software (version 6.5) with dedicated database and file server respectively. The image 24 analysis workstation used for data analysis were running ImageXpress equipped with Custom Module Editor. The DAPI channel was utilized to ensure an even cell layer without any structural defects and to identify any compounds with gross toxicity effects via visual inspection. To detect clusters of cells infected with SARS-CoV-19, we used an adaptive thresholding approach. Objects ranging from 21.9 to 250 micron with a brightness of 4500 greyscales over local background were scored as positive and the number of infected cell clusters as well as average intensity was recorded. The average number of infected cell clusters was determined for each well as mean of all sites for each well.

### Bulk RNAseq and RNAseq Analysis

Cells from a 24 well transwell ALI were collected in trizol and RNA was extracted using the Zymo Direct-zol RNA Miniprep kit (item number R2050) according to manufacturer’s instructions. The RNA concentrations were measured on a NanoDrop ND-1000 spectrophotometer. It was then analyzed by RNA-Seq at the Technology Center for Genomics and Bioinformatics (TCGB) core at UCLA.

Illumina libraries for RNA-Seq were prepared using the KAPA RNA Hyper + RiboErase HMR Kit according to manufacturer’s instructions. Briefly, the workflow consisted of depletion of rRNA by hybridization of complementary DNA oligonucleotides, followed by treatment with RNase H and DNase. Next steps included mRNA enrichment and fragmentation, first strand cDNA synthesis using random priming followed by second strand synthesis converting cDNA:RNA hybrid to double-stranded cDNA (dscDNA), and incorporating dUTP into the second cDNA strand. cDNA generation was followed by end repair to generate blunt ends, A-tailing, adaptor ligation and PCR amplification. Different adaptor barcodes were used for multiplexing samples in one lane. Sequencing was performed on Illumina HiSeq3000 for a single-end 1x50 run. Data quality check was performed using the Illumina SAV. Demultiplexing was performed with Illumina Bcl2fastq v2.19.1.403 software. RNAseq reads were mapped by STAR 2.7.9a [35] and read counts per gene were quantified using the human genome GRCh38.104. Normalization was performed using Partek Flow software [36] and read counts were normalized by CPM +1.0E-4. For comparison between samples, read counts for each gene were normalized to mock treatment or virus only treatment, respectively and the top 100 up or downregulated genes further analyzed utilizing the NIH Database for Annotation, Visualization and Integrated Discovery (DAVID) to calculate enrichment scores for functional gene clusters and calculate p-values and Benjamini-Hochberg corrected p-values.

### Z’ Calculation and Statistical Analysis

Z’ (Z-factor) for 96-well ALI was calculated following the method described by Zhang at al. [37] as described below:

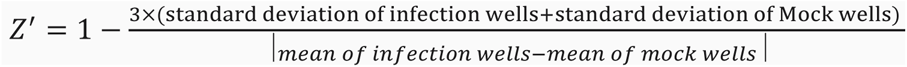

Mean and standard deviation were calculated on infection only (no drug added) and mock (no infection) wells (n>3) for each SARS-CoV-2 variant.

Student’s t-test was used to determine statistical significance of TEER value between 96-TW ALI and 24-TW ALI groups. ANOVA was used to compare the number of infection cluster across different donors and different variants. Significance was defined as p < 0.05. Statistical details of experiments can be found in the Figure legends.

### SARS-CoV-2/HeLa-ACE2 high-content screening assay

Compounds were acoustically transferred into 384-well µclear-bottom plates (Greiner, Part. No. 781090-2B). HeLa-ACE2 cells were seeded in 13 µL DMEM with 2% FBS at a density of 1.0×10^3^ cells per well. Plated cells were transported to the BSL3 facility where 13 µL of SARS-CoV-2 (strain USA-WA1/2020 propagated in Vero E6 cells) diluted in assay media was added per well at a concentration of 2.0×10^6^ PFU/mL (assay multiplicity of infection (MOI) = 0.65). Plates were incubated for 24 hours at 34°C 5% CO_2_, and then fixed with 25 µL of 8% formaldehyde for 1 hour at 34°C 5% CO_2_. Plates were washed with 1xPBS 0.05% Tween 20 in between fixation and subsequent primary and secondary antibody staining. Human polyclonal sera diluted 1:500 in Perm/Wash buffer (BD Biosciences 554723) was added to the plate and incubated at room temperature for 2 hours. Six µg/mL of goat anti-human H+L conjugated Alexa 488 (Thermo Fisher Scientific A11013) together with 8 µM of antifade-46-diamidino-2-phenylindole (DAPI; Thermo Fisher Scientific D1306) in SuperBlock T20 (PBS) buffer (Thermo Fisher Scientific 37515) was added to the plate and incubated at room temperature for 1 hour in the dark. Plates were imaged using the ImageXpress Micro Confocal High-Content Imaging System (Molecular Devices) with a 10× objective, with 4 fields imaged per well. Images were analyzed using the Multi-Wavelength Cell Scoring Application Module (MetaXpress), with DAPI staining identifying the host-cell nuclei (the total number of cells in the images) and the SARS-CoV-2 immunofluorescence signal leading to identification of infected cells.

### Uninfected host cell cytotoxicity counter screen

Compounds were acoustically transferred into 1,536-well µclear plates (Greiner Part. No. 789091). HeLa-ACE2 cells were maintained as described for the infection assay and seeded in the assay-ready plates at 400 cells/well in DMEM with 2% FBS and plates were incubated for 24 hours at 37°C 5% CO_2_. To assess cell viability, the Image-iT DEAD green reagent (Thermo Fisher) was used according to manufacturer instructions. Cells were fixed with 4% paraformaldehyde, and counterstained with DAPI. Fixed cells were imaged using the ImageXpress Micro Confocal High-Content Imaging System (Molecular Devices) with a 10× objective, and total live cells per well quantified in the acquired images using the Live Dead Application Module (MetaXpress).

### Data analysis

Image analysis was carried out with MetaXpress (version 6.5.4.532). Primary in vitro screen and the host cell cytotoxicity counter screen data were uploaded to Genedata Screener, Version 16.0.3- Standard. Data were normalized to neutral (DMSO) minus inhibitor controls (2.5 µM remdesivir for antiviral effect and 10 µM puromycin for infected host cell toxicity). For the uninfected host cell cytotoxicity counter screen 40 µM puromycin (Sigma) was used as the positive control. For dose response experiments compounds were tested in technical triplicates on different assay plates and dose curves were fitted with the four parameter Hill Equation. Technical replicate data were analyzed using median condensing.

### SARS-CoV-2/Calu-3 high-content screening assay

Compounds are acoustically transferred into 384-well µclear-bottom plates (Greiner, Part. No. 781090-2B) before seeding Calu-3 cells in assay media (MEM with 2% FBS) at a density of 5,000 cells per 20 μL per well. The plated cells are transported to the BSL3 facility where SARS-CoV- 2 (strain USA-WA1/2020 propagated in Vero E6 cells) diluted in assay media is added at an MOI between 0.75 and 1 to achieve ∼30 – 60% infected cells. Plates are incubated for 48 h at 34°C 5% CO2, and then fixed with a final concentration of 4% formaldehyde. Fixed cells are stained with human polyclonal sera as the primary antibody, goat anti-human H+L conjugated Alexa 488 (Thermo Fisher Scientific A11013) as the secondary antibody, and antifade-46-diamidino-2-phenylindole (DAPI; Thermo Fisher Scientific D1306) to stain DNA, with PBS 0.05% Tween 20 washes in between fixation and subsequent primary and secondary antibody staining.

Plates are imaged using the ImageXpress Micro Confocal High-Content Imaging System (Molecular Devices) with a 10× objective, with 4 fields imaged per well. Images are analyzed using the Multi-Wavelength Cell Scoring Application Module (MetaXpress), with DAPI staining identifying the host-cell nuclei (the total number of cells in the images) and the SARS-CoV-2 immunofluorescence signal leading to identification of infected cells.

### Uninfected host cell cytotoxicity counter screen

Compounds are acoustically transferred into 1,536-well plates (Corning No. 9006BC) before seeding Calu-3 cells in assay media (MEM with 2% FBS) at a density of 600 cells per 5 μL per well. Plates are incubated for 48 hours at 37°C 5% CO2. To assess cell viability, 2 μL of 50% Cell-Titer Glo (Promega No G7573) diluted in water is added to the cells and luminescence measured on an EnVision Plate Reader (Perkin Elmer)

### Data analysis

Data from the SARS-CoV-2 antiviral assay and host cell cytotoxicity counter screen are uploaded to Genedata Screener, Version 16.0. For the SARS-CoV-2 antiviral readout, the % CoV-2 positive cells are normalized to neutral (DMSO) minus inhibitor controls (10 µM remdesivir). For the cell count readout, the total cells are normalized to the stimulator (10 µM remdesivir) minus neutral control (DMSO). The uninfected host cell cytotoxicity counter screen is normalized to neutral (DMSO) minus inhibitor control (30 µM puromycin). For dose response experiments, compounds are tested in technical triplicates on different assay plates and dose curves are fitted with the four parameter Hill Equation. Curves are fitted as either increasing or decreasing and noted as such in the data output. This is of particular note for the cell count readout from the SARS-CoV-2 infection assay which captures both an antiviral effect, protection from virus-induced cell death (increasing), and cellular toxicity (decreasing).

### Broadband antiviral profiling

Broadband antiviral profiling was performed at IBT Bioservices using the following strains: Adenovirus 5, Chikungunya virus 181/25, Dengue virus serotype 2 D2Y98P, Herpes simplex virus 1 MacIntyre, Herpes simplex virus 2 MS, Influenza virus H1N1 A/California/07/09, Zika Virus FSS13025. 10,000 Vero cells were seeded per well in 96-well flat bottom tissue culture plates in growth medium (MEM supplemented with 10% HI-FBS, P/S and L-Gln) and incubated overnight at 37°C and 5%CO_2_. Eight 5-fold serial dilutions of each TA was prepared in serum-free medium at two times (2X) the final intended concentration; for INFV ONLY, 1 µg/mL of TPCK trypsin was added to the medium. Growth medium from the 96-well plates were then removed; 50 µL fresh serum-free medium was added to each well followed by 50 µL of each TA dilutions in triplicates; cells only and virus only wells received 100 µl of medium only. Plates were incubated at 35°C (for INFV ONLY) or 37°C and 5% CO_2_ for 60-minutes +/- 5 minutes.

Each virus strain was prepared in serum-free medium at a specific MOI of 0.01. ; for INFV ONLY, 1 µg/mL of TPCK trypsin was added to the medium. 100 µl of virus inoculum was then mixed to 100 µl of each TA concentrations, including virus only wells; 100 µl of medium only was also added to cells only wells for a final 200 µl per well. Plates were incubated at 35°C (for INFV ONLY) or 37°C and 5% CO2 for different number of days before cells fixation and staining.

After incubation, media was removed, and cells were fixed with cold 80%/20% (v/v) ethanol/methanol and incubated for 20 minutes at -20°C. Cells were washed 3x with 1xDPBS and 200 µL of the diluted anti-virus antibody (1:2000 in blocking buffer) was added to each well for overnight incubation at 4°C. Primary antibody was removed, and plates were washed three times with 1X DPBS. Common secondary antibody, Goat Anti-Mouse IgG (H+L)-HRP Conjugate diluted 1:2,000 in blocking buffer was then added at 200 µL/well and plates were incubated for 1- hour at room temperature. After removing the secondary antibody, plates were washed three times with 1X DPBS. TMB substrate (150 µL/well) was added for approximately 10 minutes (or until virus-only wells were intensely stained while cells-only wells were still weak in signal). Stop solution was then added to each well and optical density was read at 450nm.

After incubation, media was removed and cells were fixed and stained with 0.1% crystal violet solution (0.05% methanol, 50% Glutaric dialdehyde in 1xDPBS) for 60-minutes +/- 5-minutes at room temperature. Cells were washed 3x with 1xDPBS and after removing all liquid, returned plates were air dried at room temperature. Optical density was read at 570nm for each well.

Similar cell and TA preparation was done in parallel for this assay to mimic as closely as possible the antiviral assay. Virus inoculum was replaced by serum-free medium only, and for INFV ONLY, 1 µg/mL of TPCK trypsin was added to the medium. To cover all the different incubation times with only 2 conditions, plates were incubated at 35°C (for INFV ONLY) or 37°C and 5% CO_2_ for 3 and 5 days.

All media was removed and 100µL of DPBS was added to each to each, following by 100 µL of CellTiter-Glo® reagent. Plates were gently shaken for 2-minutes and incubated at RT for 10- minutes before determining luminescence.

All data were imported into Excel. The XLfit 5 plug-in was used with fit# 205 (Levenberg-Marquardt algorithm) to determine the 50 percent inhibition concentration (IC50) and the 50 percent cytotoxicity concentration (CC_50_).

### In vivo mouse studies of SARS-CoV-2 infection

In vivo experiments were performed at Vibiosphen. At D0, the mice (B6.Cg-Tg(K18- ACE2)2Prlmn/J males, 7-week-old) were pre-treated by oral gavage 1 hr before infection (100mg/kg compound in 0.5% Methocel/ 0.25% Tween-20/ water). At D0+1hr, the mice were infected with 25 µL of DMEM containing SARS-Cov-2 (1 x 104 PFU/mouse) through the intranasal route. From D0 to D5 mice were treated by oral gavage twice daily. The pause between treatments was set to 12h. At D1, D3 and D6, 5 mice from each group were euthanized by cervical dislocation for collection of lung tissue and virus quantification.

Lung tissues (up to 30 mg) were homogenized in RLT buffer and RNA was extracted using the RNeasy kit (Qiagen) according to the manufacturer’s instructions. Viral RNA was detected by qRT-PCR as described in (Corman et al., 2020) [38]:

A 25 μL reaction will contain 5 μL of RNA, 12.5 μL of 2 × reaction buffer provided with the Superscript III one step RT-PCR system with Platinum Taq Polymerase (Invitrogen, Darmstadt, Germany; containing 0.4 mM of each deoxyribont triphosphates (dNTP) and 3.2 mM magnesium sulphate), 1 μL of reverse transcriptase/Taq mixture from the kit, 0.4 μL of a 50 mM magnesium sulphate solution (Invitrogen), and 1 μg of nonacetylated bovine serum albumin (Roche). All oligonucleotides were synthesized and provided by Tib-Molbiol (Berlin, Germany). Thermal cycling was performed at 55 °C for 10 min for reverse transcription, followed by 95 °C for 3 min and then 45 cycles of 95 °C for 15 s, 58 °C for 30 s. Participating laboratories used either Roche Light Cycler 480II or Applied Biosystems ViiA7 instruments (Applied Biosystems, Hong Kong, China). Ten-fold dilutions of SARS-CoV-2 standards with known copy numbers were used to construct a standard curve.

---

**Supplementary Table 1.** List of donor demographics, drug libraries and SARS-CoV-2 variants.

| Donor | Demographics | Drug library | Variant tested |
| --- | --- | --- | --- |
| DL1 | 70, M, B, Non-NS, A | LOPAC | Wuhan |
| DL2 | 66, M, C, S, A | LOPAC, New Prestwick | Wuhan, Beta |
| DL3 | 59, M, C, NS, A | LOPAC, New Prestwick | Wuhan, Beta, Delta |
| DL4 | 79, F, C, S, A | NKIL | Wuhan, Delta |
| DL5 | 55, F, C, S, A | NKIL | Delta, Omicron |
| DL6 | 51, M, C, NS, NA | All previous hits | Wuhan, Omicron |
| DL7 | 67, F, C, NS, A | All previous hits | Wuhan, Omicron |
| DL8 | 45, F, C, S, A | All previous hits | Wuhan, Omicron |
| Pooled DL6-8 | As above | Top hits+Merck | Omicron, Nanoluc |

M=male, F=female, B=Black, C=Caucasian, S=smoker, A=alcohol use, NS=nonsmoker, NA = no alcohol use

